# Unsupervised identification of internal perceptual states influencing psychomotor performance

**DOI:** 10.1101/2023.09.08.556817

**Authors:** Ozan Vardal, Theodoros Karapanagiotidis, Tom Stafford, Anders Drachen, Alex R. Wade

## Abstract

When humans perform repetitive tasks over long periods, their performance is not constant. People may drift in and out of states that might be loosely categorised as engagement, disengagement or ‘flow’ and these will be reflected in multiple aspects of their performance (for example, reaction time, accuracy, criteria shifts and potentially longer-term strategy) but until recently it has been challenging to relate these behavioural states to the underlying neural mechanisms that generate them. Here, we took Magnetoencephalograpy recordings of participants performing an engaging task that required rapid, strategic behavioural responses. In this way we acquired both high density neural data and contemporaneous, dense behavioural data. Specifically, participants played a laboratory version of Tetris which collects detailed recordings of player input and game-state throughout performance. We asked whether it was possible to infer the presence of distinct behavioural states from the behavioural data and, if so, whether these states would have distinct neural correlates. We used hidden Markov modelling to segment the behavioural time series into states with unique behavioural signatures, finding that we could identify three distinct and robust behavioural states. We then computed occipital alpha power across each state. These within-participant differences in alpha power were statistically significant, suggesting that individuals shift between behaviourally and neurally distinct states during complex performance, and that visuo-spatial attention change across these states.

## 1 Introduction

Digital games are a promising paradigm for research into human cognition. In recent years, researchers have used telemetry data recorded in commercial games to investigate theories of motor chunking [1, 2], ageing [3, 4], and sleep consolidation [5], among some examples of problem domains. On the other end of the methodological spectrum, games that have been tailor-made for laboratory research have been used to further our understanding of neural plasticity [6, 7], skill transfer effects [8, 9], and have been used to test cognitive architectures that model human learning as a whole [10].

One problem associated with games in research is the use of total or end-game scores to compare the aggregated performance of groups of individuals. Unfortunately for investigators interested in sterile research environments, games are frequently complex and designed to be engaging, often meaning that players encounter variations of problem spaces across separate interactions with a given game. This makes the analysis of performance using total scores difficult, as they may mask underlying factors that can vary across trials and sessions, for instance, changes in player behaviour as a response to novel situations in the game. A proposed solution is that researchers look past total scores by interrogating variables describing *components* of performance, such as patterns of players’ control inputs and decisions [11, 12, 13].

This is a particular advantage afforded by digital games as experimental tasks, as their programming often allows researchers to extract high-density behavioural data describing multiple aspects of performance.

A related, broader problem, and one that is potentially exacerbated by the variability in problem spaces inherent in digital games, is that individuals may alternate between periods of good and bad performance within single sessions of play despite proficiency in the game. In behavioural neuroscience, trial-to-trial variability is present even in low-level psychophysics tasks. These confounding behavioural observations have often been the subject of modelling efforts, whereby researchers seek to explain fluctuations in observed behaviour in terms of changes in latent cognitive factors, such as noise in perceptual systems or shifts in attention [14, 15]. In recent years, novel applications of unsupervised learning techniques have allowed researchers to statistically relate trial-to-trial variability to these unobservable cognitive factors, typically modelled as discrete shifts in internal *states* [16, 17, 18]. Inspired by these approaches, the aim of this study is to address the two problems described above through the identification of latent perceptual states in the context of human psychomotor performance. More specifically, using simultaneous, high-density behavioural and magnetoencephalography (MEG) recordings of participants playing Meta-T [19], a laboratory version of Tetris, we show that the dynamics of psychomotor performance can be modelled as observations influenced by latent perceptual states, which we relate to neural markers of attention.

A convenience assumption that is made in many cognitive experiments involving sequential measurements, is that data are independent and identically distributed samples from a shared distribution. We can consider an alternative and arguably more realistic perspective by assuming that multiple processes with distinct mechanisms contribute to recurring sequences of observations in a given data set [20]. By treating these temporally recurring patterns in time series as being influenced by distinct, unobservable processes, we can segment continuous data into patterns of observations based on inferences about underlying latent states. For example, in the context of digital games, differences in patterns of performance may arise from players consciously adopting different behavioural strategies [21], or from players shifting attention between different aspects of skill [22]. Modelling the dynamics of moment-to-moment performance in this way can allow us to make sense of fluctuations in individuals’ performance that are typically dismissed as noise, but may actually be important sources of information.

Previous studies adopting this modelling framework have shown that brain states during waking behaviour shape the dynamics of cortical activity, stimulus-response and task performance in different animals and in recent cases have been able to describe behaviour with striking accuracy [23, 24, 25, 17]. For instance, researchers investigating the acoustic courtship behaviours of fruit flies were able to precisely predict distinct patterns of song behaviour by statistically inferring latent states from flies’ movement data, capturing 84.6% of all remaining song patterning information that previous models lacking a latent state component could not explain [17]. Accurate segmentation of the latent state sequence allowed detailed description of the flies’ sensorimotor-strategies corresponding to each state and, following an optogenetics component of the study, identification of the neurons responsible for switching between states. Similarly, latent state models of rodent decision-making can accurately predict choice strategies corresponding to states of optimal engagement versus bias [26, 18], permitting reliable detection of blocks of trials with heterogeneous error-rates, as opposed to previous models that would assume errors are scattered throughout all trials in a session with equal probability.

While varying in scope and problem domain, common to some of these studies is their use of Hidden Markov Models (HMM), which model observable processes in terms of an underlying sequence of unobservable (i.e., hidden) states that transition with fixed probabilities. In general, the typical modelling pipeline involves specification of the number of states that are assumed to influence the process as well as the probabilities of the model initializing each state, following which the parameters of the model are estimated via maximum likelihood estimation. As described previously, successful validation of HMMs in cognitive task environments allows post-hoc relation of observable behavioural dynamics to underlying brain states, resulting in rich descriptions of moment-to-moment performance and cognition. These can exist at the group level but also the individual level, for instance by analysing how much time individual participants spend in each state and how often they transition between states [27].

Depending on the objectives of modelling, the specification of the states can take on different forms. In the examples outlined above, researchers specified a distinct generalized linear model (GLM) for each state that acted as a psychometric function mapping stimulus to sensorimotor response [17, 18]. This approach paired the HMM with a previously tested GLM with proven application in tasks with discrete outputs. A similar usage tested stagewise models of human skill acquisition by pairing each latent state (i.e., stage of learning) with a different speedup function describing participants’ response latencies in a novel arithmetic task [28]. Other investigations of latent states in humans have included the identification of brain states during wakeful rest or motor task performance by fitting HMMs to electrophysiological time series [27, 23, 29]. As such, studies have demonstrated that HMMs provide a flexible and task-agnostic framework for segmenting behavioural or neural time series into meaningful state-dependent epochs.

### 1.1 Neural correlates of internal states

Thus far, we have highlighted how trial-to-trial variability in psychomotor data obtained through digital games (and in psychological data sets in general) remains unaccounted for in many studies, despite the burgeoning use of digital games as paradigms to investigate psychomotor learning and other aspects of cognition. We have also provided an overview of how sequence classification of high-density behavioural or neural time series, in particular through the use of HMMs, can help researchers to make sense of trial-to-trial variability across various task environments by identifying latent states that subjects shift between as they engage in a task. Accordingly, we continue by considering how latent state identification through this approach may be applied to high-density psychomotor data. In particular, we ask the question: what might states identified through such means represent in terms of their underlying neural and cognitive dynamics?

One aspect of neural activity that can inform us about internal states is endogenous rhythms. These refer to the cyclical patterns of neural oscillations that occur naturally in the brain (independent of external stimuli) as neurons settle into stable firing rates. Endogenous rhythms can be observed at various levels of neural organization, from individual neurons to large-scale networks, and can be characterized by their frequency, amplitude, and phase. Some examples include oscillations in the *alpha* (8-12Hz), *gamma* (25-80Hz), and *theta* (4-8Hz) frequency bands, which have been linked to various cognitive functions such as attention [30, 31, 32], learning [33, 34], and memory [35, 36]. While endogenous rhythms are also associated with coordination and communication between different brain regions [37], we focus presently on the cognitive correlates of rhythms within individual brain regions.

The alpha rhythm was the first human brain rhythm to be detected in the human brain, and is easily measurable across the cortex using electrophysiological methods [38]. Despite ongoing conflicts regarding the underlying mechanisms, there is a well-established relationship between occipital alpha and visual spatial attention [39, 40]. More specifically, direction of the attentional “spotlight” from one location in the visual field to another in the absence of eye movement has been shown to correlate with modulations in the amplitude of alpha rhythm in both the parietal and primary occipital cortices [41, 42]. Alpha can be robustly detected in occipital regions, where it is thought to be associated with visual attention through the suppression of task irrelevant information [43, 44]. Additionally, studies have demonstrated that neural activity in this area is modulated by attention even when visual stimuli are not present [45, 46].

### 1.2 Aims and approach

Crucially, the approach described here depends on multivariate input data, such as pupillometry and movement in addition to task outcome. In contrast, many studies of human performance that use commercial games are limited to analyses of outcome scores, such as match wins or number of points scored in a round. We bridged this gap by using a previously tested version of a commercial game (Tetris) that has been adapted for use in the laboratory. This version of Tetris records control inputs at every stage of the game, and outputs logs of variables that are of interest to cognitive researchers, such as response latency and motor efficiency. Our aim was to identify latent states (e.g., states of high versus low engagement), to characterise recurring patterns of performance during gaming. Using MEG, we then investigated cortical activity relating to attention in each state. To our knowledge, this is the first investigation of latent states to be conducted in an ecologically valid context of human psychomotor performance.

To validate the use of this task for studying behavioural states, we first performed a purely behavioural experiment (E1) using secondary experimental data from an independent lab. Data from E1 were used to decompose game-state and behavioural logs into orthogonal components of Tetris performance, and to test the capacity of these performance components to distinguish between periods of good and bad performance. After validating our dimensionality reduction pipeline, we performed a new experiment (E2) inside the MEG scanner where we recorded neuronal activity while subjects played the game. We then used the dimensionality reduction pipeline from E1 to extract performance components from E2 behavioural data. This was followed by analysis of synchronised behavioural and neuronal data from E2.

## 2 Methods

We used a Python 3.8 [47] environment for all preprocessing and analysis. Data munging and preprocessing were performed using pandas [48] and [49] and we used matplotlib and seaborn for visualisation. Additional packages for corresponding analysis techniques are detailed below under Results. All analysis pipelines and supporting software are publicly available at *https://github.com/ozvar/tetrisMEG*, together with details regarding requisite software dependencies. Figure 1 provides an overview of the experimental and analysis pipelines.

**Figure 1:**
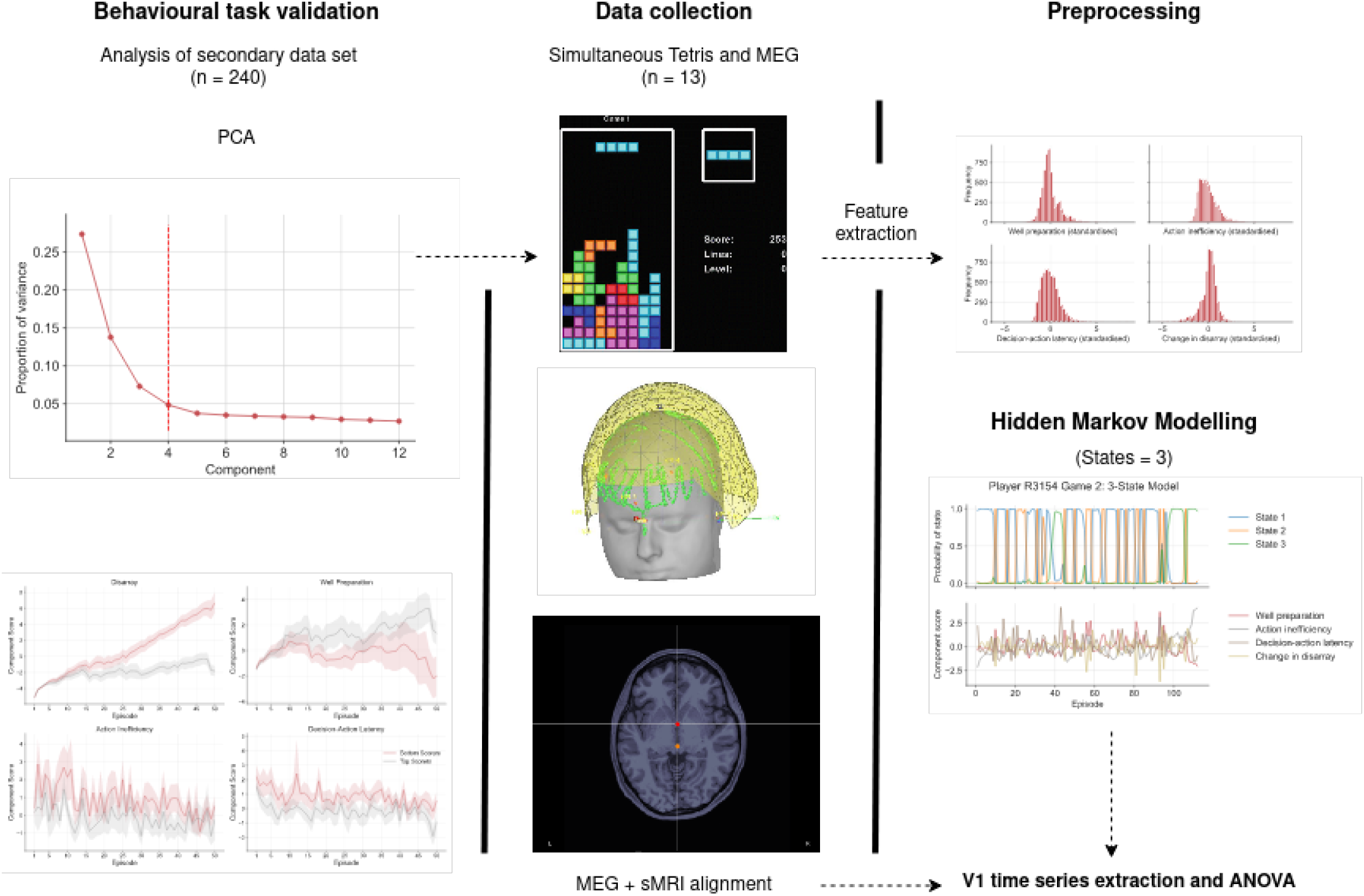
Overview of experimental and analysis pipelines. **Left column:** We first validated our feature extraction protocol on an archival data set of 240 participants, confirming that Meta-T reliably produces meaningful behavioural information relating to motor planning and execution. **Middle column:** We then collected simultaneous behavioural and MEG data using the same task, instructing participants to play Meta-T under the scanner. Structural MRI scans were also taken to permit the estimation of MEG sources. **Right column:** PCA weights extracted from the archival data set (left column) were then applied to the behavioural data obtained in our experiment (middle column). After fitting a three-state HMM to the resulting principal component scores, we aligned MEG and sMRI data for each participant, and computed alpha power in the visual cortex across each HMM state.

### 2.1 Task validation

The behavioural task was a pygame implementation of Tetris called “Meta-T”, developed by Lindstedt and colleagues for the purpose of studying human expertise and learning [19]. Meta-T is a near-identical representation of the original NES Tetris, with the exception of minor visual differences relating to the use of Python as the development language.. Importantly, Meta-T possesses several additional features that make it suitable for cognitive science, and has been used as a task environment in several published studies on human and machine expertise [50, 51, 52, 53]. Firstly, Meta-T outputs several data files at the end of each session that, in addition to detailing the participant’s ID and other session-specific information, include a log of post-game summary statistics for each game, a log of game-state (e.g., pile structure) and behavioural (e.g., action latencies) information describing performance for each “episode” of play (i.e., the time between a tetromino appearing to the time at which it is placed), as well as a complete log containing key-press information at the millisecond level (See [19] for an exhaustive description of logged variables). Secondly, researchers can modify game parameters such as the screen size, game length, or difficulty curve, by editing the default configuration file. In doing so, researchers can constrain participant behaviour to bespoke experimental conditions according to the requirements of their research question.

We used a secondary, experimental data set of Meta-T gameplay made public by Lindstedt and colleagues through the Open Science Framework (https://osf.io/78ebg/). We describe the data set here following the original experimenters’ reports as well as our own examination of its contents. The data comprise one hour of Meta-T performance from each of 240 participants, collected under laboratory conditions. Meta-T was configured to resemble “Classic Tetris” (and verified as a faithful representation thereof by a software expert affiliated with the Classic Tetris World Championship). Each participant was seated in front of a computer and instructed to play 50 minutes of Meta-T with a provided Nintendo Entertainment System controller. Players repeatedly played successive games of Tetris until 50 minutes had elapsed restarting games upon failure. The data set comprise three log files, each detailing all 240 participants’ task engagement at three different levels of time: one describing behaviour at the time of each button input, one describing behaviour in the time spanning the appearance to dropping of each tetromino, and one summarising behaviour at the level of the entire match. We concentrated our analyses on logs of tetromino drops at each match, as these provided the highest density of measures across all log files. As mentioned previously, Meta-T captures a wealth of information throughout gameplay. Each row in the episodic log file details over 60 variables for the current tetromino drop, including:

1. Features summarising the session (e.g., participant ID, game number, timestamp),
2. Game state features relating to the tetromino (current and next tetromino, current tetromino position),
3. Game state features describing the pile (e.g., height, circumference, number of unplayable cells)
4. Motor execution features (e.g., number of control inputs, latency before and between actions),
5. Features describing tetromino placement (e.g., number of lines cleared, landing height).

We refer the reader to Lindstedt and colleagues [53] for an exhaustive list of all variables that are logged by the Meta-T. In the next section, we describe the variables we chose to concentrate our analysis on, which were principally those describing the pile structure and those describing motor execution.

#### 2.1.1 Principal component analysis

Our first aim was to reduce the data set into a subset of variables relating to orthogonal aspects of performance in Tetris. To this end, we performed an exploratory principal component analysis (PCA) using the sklearn package [54]. We concentrated this analysis on episodic logs describing behaviour and game-state at the level of each tetromino drop, as these logs provided the greatest breadth of information relating to moment-to-moment performance. For each game that was played, Meta-T produced one such log file as a .tsv spreadsheet. Each row in these files corresponded to one tetromino drop, describing the input behaviours from the moment the tetromino appeared to when it was dropped, changes to the game-state following the tetromino drop, as well as summary variables describing participant and session related information.

We first inspected the data for extreme outliers and other anomalies, removing two players from the data set who never made it past level 0 during gameplay. We then subset the data, removing all variables describing the session (e.g., participant ID, time stamp) or game-state (e.g., current tetromino, tetromino orientation), effectively retaining only those variables relating to performance for dimensionality reduction (see Table 2 for a list of variables retained for PCA). PCA was then performed on this trimmed data set, initially with an unconstrained number of components.

To identify the optimal number of components to retain, we produced a line plot of the amount of variance explained by each successive component (Figure 9). Identifying the point of maximum curvature in this plot indicated four components as being the optimal number to retain, as adding more would have had a limited impact on the explained variance. We provided meaningful labels to each of these components, similar to Lindstedt and colleagues [53], according to the unique aspect of Tetris performance captured by each one. Together these components explained up to 53.3% of the variance in Meta-T performance. Table (2) displays the PCA loadings, describing the correlation between each variable and principal component (only correlations past 0.20 are displayed). We take a moment here to interpret each component in detail.

1. *Disarray*. Players that fail to clear lines as their tetris pile increases in size are prone to developing an unfavourable tetris pile. Disarray captures this deficiency in pile structure, and is associated with unplayable holes and jaggedness of the pile, as well as overall pile height.
2. *Well preparation*. Achieving a high score in Tetris requires capitalising on opportunities to score bonus points, typically by clearing multiple lines with a single tetromino. Well preparation relates to the forward planning required to achieve multiple line clears, such as by reserving a single, deep well and maintaining a neat pile structure.
3. *Action inefficiency*. Action inefficiency captures inputs (e.g., rotations, translations) that are made in excess of the minimum number of inputs required to place a tetromino at its final destination. This may relate to poor motor execution and planning.
4. *Decision-action latency*. This component corresponded to the initial lag and average lag between actions associated with each tetromino placement. It also corresponded to the local quality of placement for each tetromino (i.e., the reduction in pile height caused by placement and amount of contact with tetrominoes in the pile). Taken together we view this component as capturing both the speed and quality of decision-making as it relates to identifying optimal tetromino placement.

Taken together, the PCA four components that we believe captures distinct aspects of Tetris, each of which may act as proxy measures of cognitive-behavioural aspects of psychomotor performance. Our results also resemble dimensionality reduction performed by Lindstedt and colleagues, who retained three orthogonal components in their analysis using the same data set [53]. We retained one component describing (sub)optimality of the game state, one component describing strategic planning in relation to pile structure, and two components that potentially tap into aspects of executive function, such as working memory and planning.

To put the results of our PCA into the context of task performance, we visualised the averaged trajectories of each of these components across the first 50 tetromino drops of two example games from the sample (Figure 2). We chose to visualise 50 drops as over half the sample had played at least this many tetrominoes in every one of their games. Evident in this visualisation is that disarray appears to monotonically increase across both games, likely relating to increase in pile complexity as the game goes on. Well preparation also appears to trend upwards, as would be expected from players attempting to play strategically in the face of incoming tetrominoes. The final two components appear to be statistically stationary, fluctuating without any visible periodicity within these first 50 episodes.

**Figure 2:**
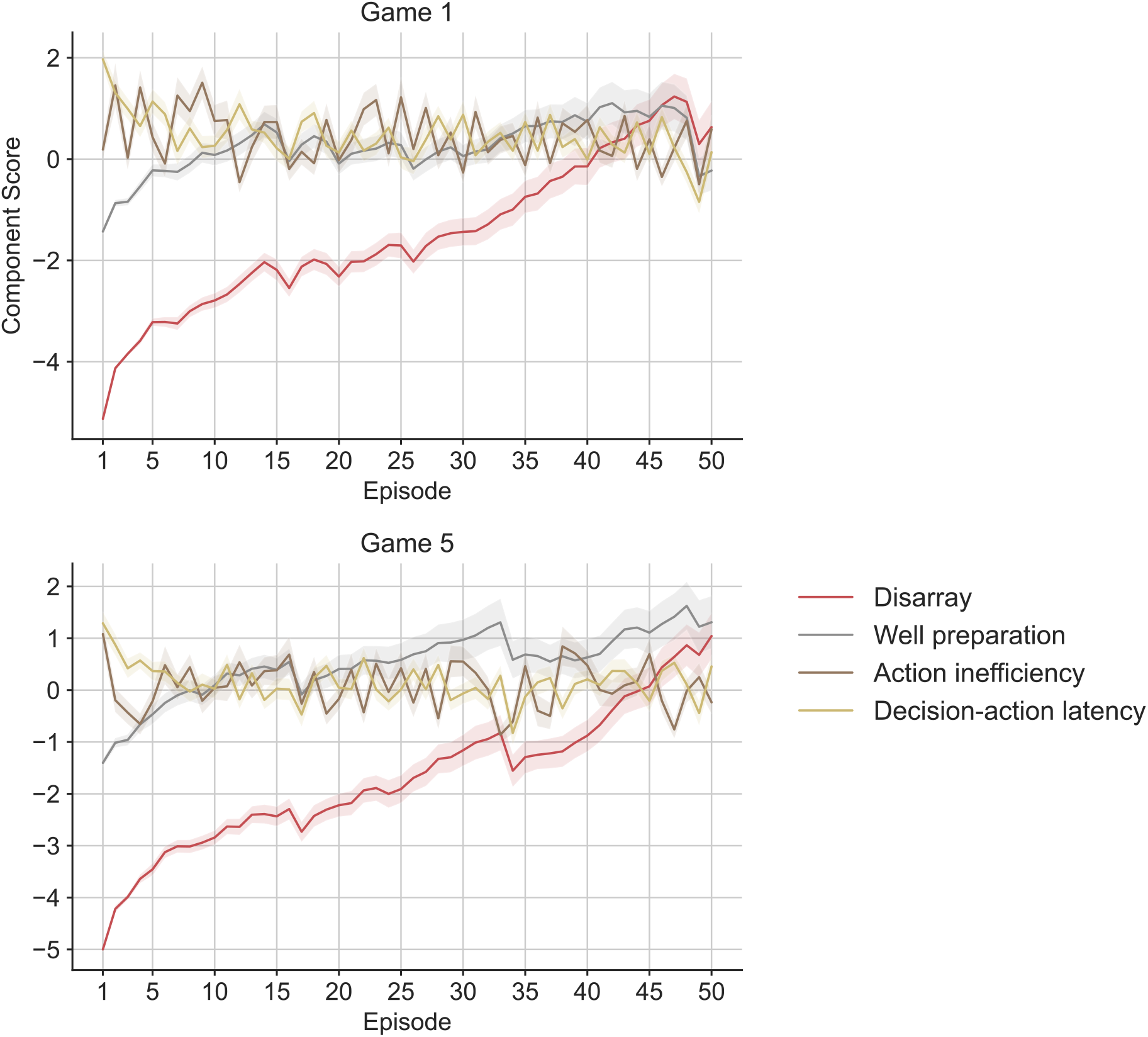
Averaged component score trajectories across two exemplar games. The top and bottom panels show mean disarray, well preparation, action efficiency, and decision-action latency from the first and fifth game across the entire sample (including only those participants who played at least five games) respectively. Shaded regions show 95% confidence intervals of the mean.

#### 2.1.2 Distinguishing top from bottom scorers

If these measures are meaningful, we would expect them to to distinguish between good and bad performances, or between heterogeneous groups that exist within our sample, such as top scoring players versus bottom scoring players. To investigate the latter idea, we first split players into a top and bottom scoring group by sorting players based on their average score on their first few games, and then taking the top and bottom quintiles respectively for each group. We then visualised, for each group separately, the averaged trajectory of each component over the first 50 episodes of the first game played (Figure 3).

**Figure 3:**
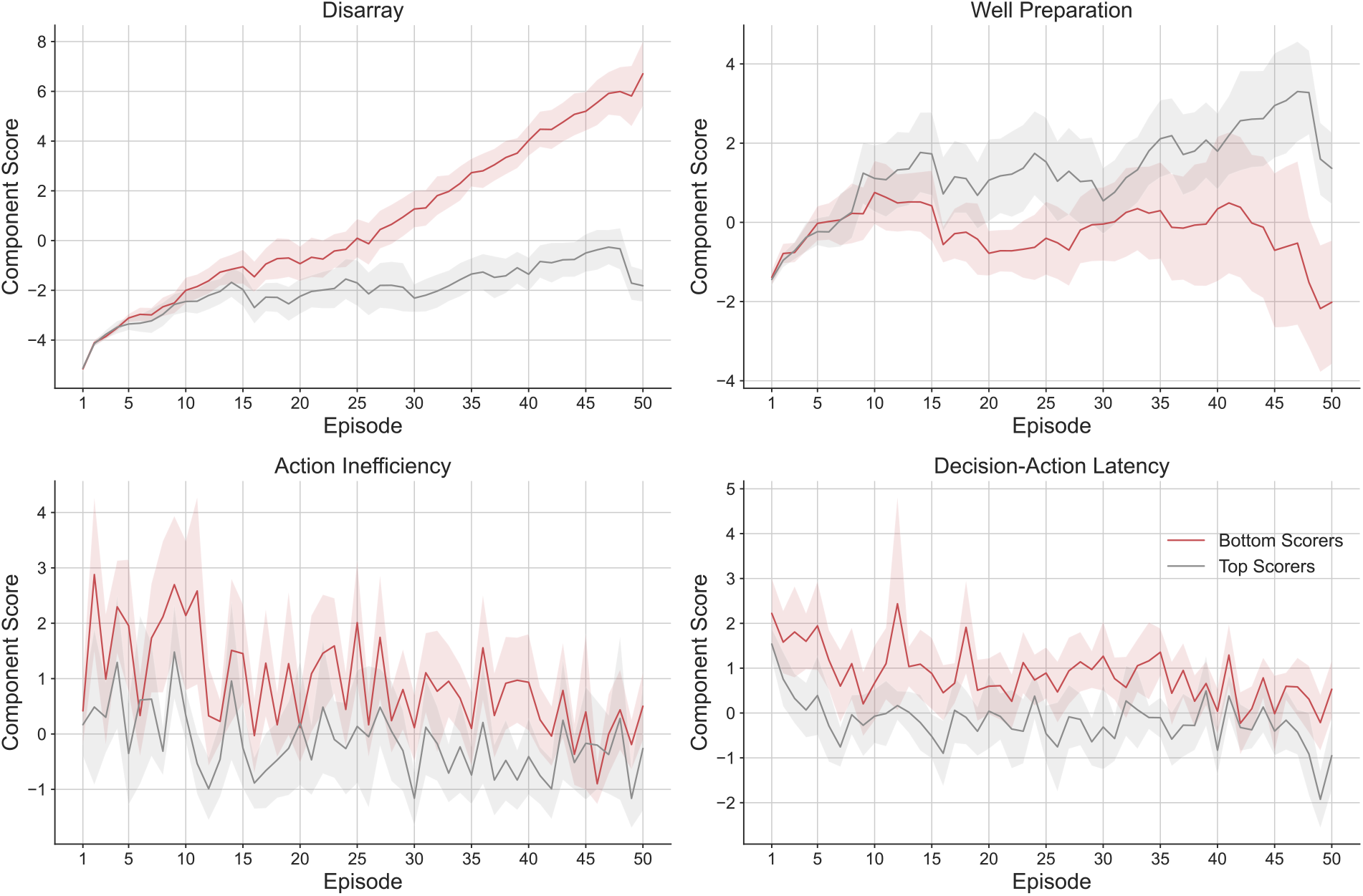
Comparison of moment-to-moment performance between top and bottom scorers. Each panel depicts the mean trajectory of the respective performance component for bottom (red line) versus top scorers (grey line) for the first 50 tetromino drops from game 1. From top left going clockwise to the bottom left panel, the subplots show trajectories for mean disarray, well preparation, decision-action latency and action inefficiency respectively. Shaded regions depict 95% confidence intervals of the mean.

It is evident across every panel that top scoring players differ significantly in mean respective component score across time. As a reminder, participants are exposed to identical tetromino sequences for each successive game that is played. Thus, it is striking that while the peaks and troughs in action inefficiency and decision-action latency appear similar between top and bottom scorers, the top scorers appear to be more efficient and quick in their gameplay at almost every tetromino drop in the game. Moreover, while disarray in both groups appears to trend upwards, the upward trend is much more aggressive in bottom scorers than in the top scoring group. Conversely, bottom scoring appears trend downwards in their well preparation while top scoring players appear to trend upwards.

We tested the statistical significance of visible trends in our plots by conducting a mixed ANOVA of each performance component, with scoring decile as the between subjects factor, and tetromino drop as the within-subjects factor. Between-subjects effects for disarray [F(1, 34) = 81.93, *p* < 0.001, partial *η*^2^ = 0.71], well preparation [F(1, 34) = 10.90, *p* = 0.002, partial *η*^2^ = 0.2427], action inefficiency [F(1, 34) = 26.38, *p* < 0.001, partial *η*^2^ = 0.44], and decision-action latency [F(1, 34) = 21.75, *p* < 0.001, partial *η*^2^ = 0.39] were all statistically significant.

Additionally, interaction effects between scoring decile and tetromino drop were statistically significant for disarray [F(1, 34) = 36.79, *p* < 0.001, partial *η*^2^ = 0.52], well preparation [F(1, 34) = 3.84, *p* < 0.001, partial *η*^2^ = 0.11], and action inefficiency [F(1, 34) = 1.42, *p* = 0.30, partial *η*^2^ = 0.04], but not for decision-action latency [F(1, 34) = 1.09, *p* = 0.31], suggesting that between-groups differences in the former three components grew statistically significantly more pronounced as games went on.

### 2.2 Participants

15 healthy, right-handed participants were recruited through the York Neuroimaging Centre (YNiC) participant pool (United Kingdom). All participants provided informed consent, and the study was approved by the York Neuroimaging Centre ethics committee. All participants were familiar with playing Tetris, and provided a self-report of their proficiency on a 5 point Likert scale (*M* =3.08, *SD*=1.04), as well as their proficiency in digital games in general (*M* =3.38, *SD*=1.19). Data from 2 participants were excluded from analysis due to poor MEG data quality, resulting in a final sample of *n* = 13 participants (4 female, *M* _age_ = 33, *SD*_age_ = 11.31).

### 2.3 Behavioural data acquisition

Participants played an updated version of the pygame implementation of Tetris used in E1 (Figure 4). Several adaptations were made to facilitate a Python3 environment as well as to enable synchronisation with MEG recording. For the present study, we configured Meta-T to run in a full-screen environment without in-game music. We also fixed the set of numerical seeds determining the tetromino sequence in each game, such that each subsequent game had a different sequence of tetrominoes, but the variations in tetromino sequence were the same across all participants. Games had an indefinite length (i.e., players played each game until loss) and all other options were left at the default values for Classic Tetris. The original stimulus code was further adapted to accommodate a fibre optic response interface (Cambridge Research Systems 905 package) connecting between the stimulus computer and MEG scanner, which allowed participants to use a non-electronic, non-magnetic five button response pad to play Meta-T without adding additional noise to the scans. We configured Meta-T to send triggers to the MEG record via this interface upon the occurrence of salient events. These included button inputs, tetromino appearances and drops, line clears, as well as game start and game end.

**Figure 4:**
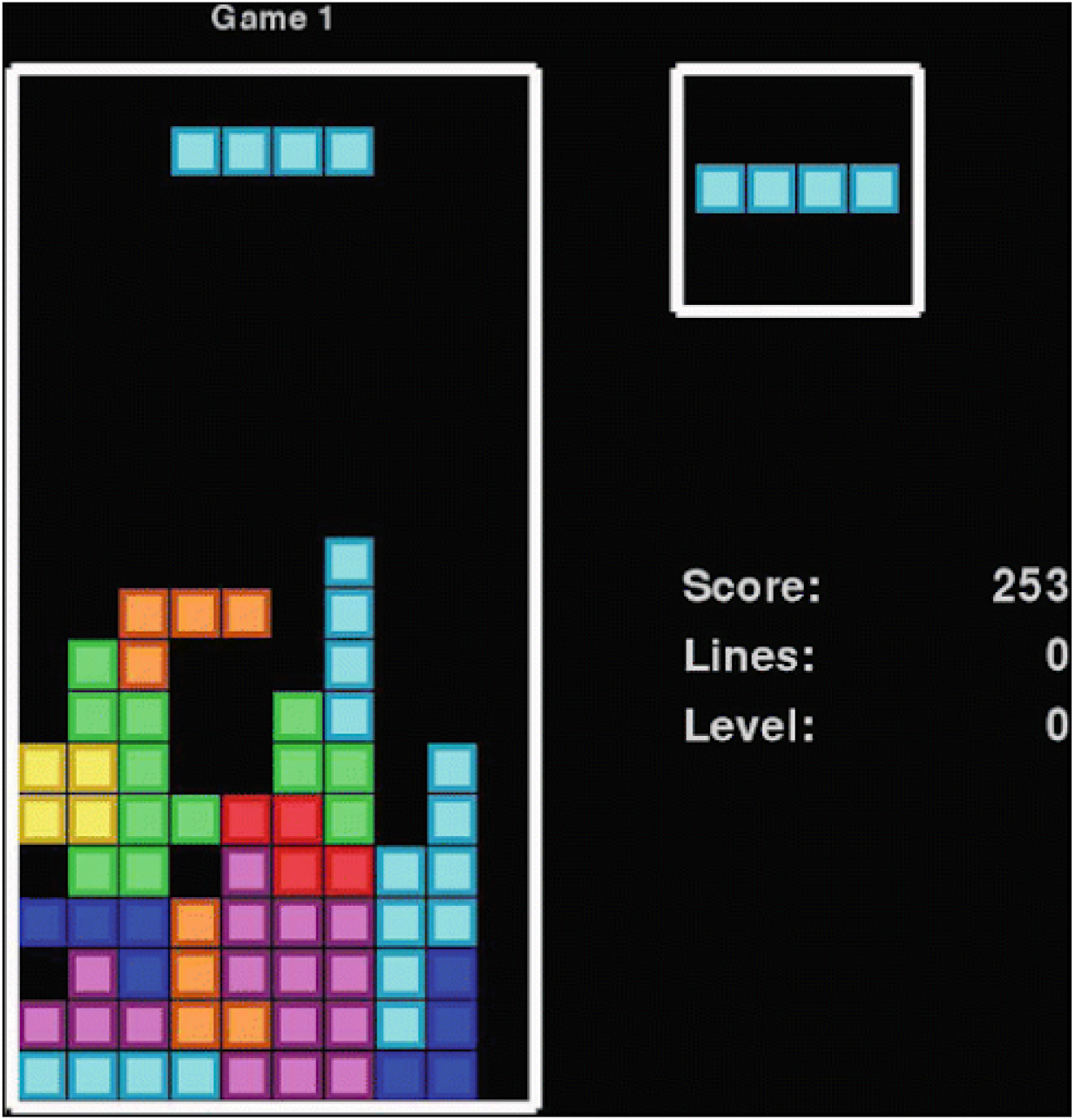
Depiction of Meta-T user interface. The left side of the screen shows the current board, including the tetromino that is currently being controlled above it. The top right side of the screen shows the *next* tetromino that will be played after the current one is dropped. The player is also presented with the current game number, their current score, number of cleared lines, and the current level. Image taken from Lindstedt and Gray (2019).

### 2.4 Magnetoencephalograpy (MEG)

MEG scanning was conducted using a 4D Neuroimaging Magnes WH3600 scanner (248 channels + 23 reference channels) at YNiC and data were acquired at a sampling rate of 500 Hz and then downsampled to 200 Hz. Prior to scanning, five feducial head-coils were attached to each participant’s head with hypoallergenic tape. Facial landmarks (nasion, left and right preauricular) and head shape were then recorded using a Polhemus Tastrak 3D digitizer. To assess head movement inside the scanner helmet, we measured the position of the head-coils before and after every scan, and then compared these measurements to the spatial relation between head-coils recorded outside of the scanner. Movement *<* 0.5cm was our acceptance threshold for head movement, beyond which we reran our coil-on-head (CoH) scan to confirm any discrepancies in coil position and to subsequently recalculate coil positions using non-displaced coils.

After being briefed and prepared for scanning, each participant was given some time to practice playing Meta-T in the scanner until they reported feeling well-adjusted to the button inputs, during which time the data acquisition software was configured for scanning. Participants played Meta-T in a seated position and were instructed to play until they lost, while keeping their head as still as possible. Each scan was initiated five seconds before the start of each game and scan duration varied for each participant depending on their performance across games (i.e., better performance resulted in longer games). As described above, each scan and concurrent game was preceded and followed by a CoH scan, allowing us to assess head movement while the participant took a short break. Each participant typically played two or three games, resulting in an average acquisition duration of *M* = 7.84 minutes per game (*SD* = 2.88) and an average total acquisition duration of *M* = 21.55 minutes per participant (*SD* = 4.54).

### 2.5 Structural magnetic resonance imaging (MRI)

To estimate the neural sources of our MEG recordings, MEG data were co-registered with high-resolution structural MRI images. T1-weighted structural MRI scans were acquired for each participant using a Siemens Prisma 3T MRI scanner, and the Freesurfer pipeline [55, 56] was used to perform image segmentation and cortical reconstruction. Head surfaces digitized during MEG preparation were then aligned with reconstructed MRI images based on the aforementioned fiducial landmarks.

### 2.6 Analysis

#### 2.6.1 Dimensionality reduction of behavioural data

We transformed our behavioural data set using weights from the PCA that was performed on the previous, secondary data set. We used the weights from the model to transform the behavioural data in the current experiment, producing for each participant an additional time series of four components relating to the aspects of Tetris performance described above. As we were primarily interested in analysing moment-to-moment changes in performance, we used the mathematical difference of disarray in lieu of the default disarray value, as the former assesses the impact of each tetromino drop on pile structure while the latter provides a status desription of the same. The scores of each component were then standardised to permit comparison between components with different scales.

#### 2.6.2 Hidden Markov Models

We used the Python hmmlearn package (an open source module with an API similar to scikit-learn; hmmlearn, [57]), to fit a three state HMM to the time series of PCA-derived performance variables, where each point in the time series describes participant performance at each tetromino drop. Our model was fit to our data at the group-level (as in Karapanagiotidis et al., [29]) by concatenating the data across all of our participants and games. We fit a Gaussian HMM (the observations are assumed to be well-described by a Gaussian distribution) with a diagonal covariance matrix and a 200 iteration upper bound for training, ensuring that the Expectation Maximisation (EM) algorithm stopped either after 200 iterations or on convergence to a maximally likely solution before reaching the iteration limit. As the EM algorithm is gradient-based and may therefore converge to local optima, we ran multiple courses of model fitting with different initializations (a random initial transition matrix for the states) but otherwise identical parameters. We then chose the model with the highest log-likelihood for the remainder of our analyses. As an additional check of model robustness, we compared the log-likelihood of our true model to a randomised chancel model that we produced by fitting an HMM with identical parameters to a randomly shuffled time series of our observations. We observed a consistently higher model fit in our true model as compared to our chance model.

We chose a three state model assuming three modes of engagement with Meta-T: a default state where participants were engaged and attentive, a performant state where participants were both engaged and playing optimally, and a “panic” state involving suboptimal moves and blunders, potentially relating to inattention. States were assigned to each point in the behavioural time series using the Viterbi algorithm, which finds the most likely sequence of hidden states given the observations. It is worth noting that due to the variable amount of time taken by players to drop each tetromino, each point in the resulting HMM state time series was also of variable length.

### 2.6.3 MEG data pre-processing

Analysis and pre-processing of MEG was performed in MatLab using Brainstorm [58]. Data were first band-pass filtered between 1 and 40 Hz using a finite impulse response filter. We performed an Independent Component Analysis (using the infomax algorithm; Bell & Sejnowski, [59]) to reduce the MEG data and proceeded to identify and reject components capturing periodic physiological artefacts such as blinks and heartbeats. Raw time series for each scan were then inspected manually in epochs of 50 seconds, and any periods contaminated by additional artefacts were manually removed.

After co-registering the MEG data from each scan with the corresponding participant’s structural MRI data, we used the minimum-norm imaging function in Brainstorm to estimate the amplitude of sources across the cortical surface via minimum-norm estimation [60]. To how brain activity changed across each behavioural state, we needed to examine MEG data Extract source time series for each state, we first aligned the time series of the behavioural and MEG recordings, before segmenting the behavioural time series into bins of one second in length. For each bin, we then extracted the time series of squared amplitudes from regions of interest (ROIs) for each bin and state, parcellating ROIs from left and right hemispheres using the Brodmann atlas. We maintained a uniform length of one second for each bin to ensure that fourier transforms of each bin, computed with the Fast Fourier Transform algorithm (FFT), were equal in length. Finally, we obtained a measure of the mean amplitude for frequency bands across each state, by computing the root-mean square (RMS) of amplitude across frequencies for all bins for each given state, and each participant. Specifically, we computed the RMS of alpha rhythm (8-12Hz) in V1, and RMS of mu rhythm (8-13Hz) in M1 across each HMM state for each participant.

## 3 Results

### 3.1 SVM decoding

To assess data quality, we checked that we were able to distinguish between neural responses to button presses executed with the left versus right thumb (i.e., the buttons used to translate the tetromino left and right respectively), as this would provide us with confidence that our MEG and behavioural data were both adequately synchronised and contained meaningful information. For each participant, we trained a linear Support Vector Machine using the libsvm library [61] to decode left versus right translation inputs using the MEG time series extracted from −400ms to 400ms relative to each button press. To improve computational efficiency and signal-to-noise ratio, trials from each class (i.e., left versus right translation) were randomly assigned to 5 folds. Trials in each fold were then subaveraged, yielding a total of 5 subaveraged trials per class. Decoding was then performed on the 5 subaveraged trials following a leave-one-out cross-validation procedure, and the process was iterated 50 times. Classification accuracy was averaged across the 50 iterations for each millsecond across the trial time range, and plotted for each participant (Figure 5). We attained a classification accuracy of 100% for every participant approximately 0.1 seconds after the response.

**Figure 5:**
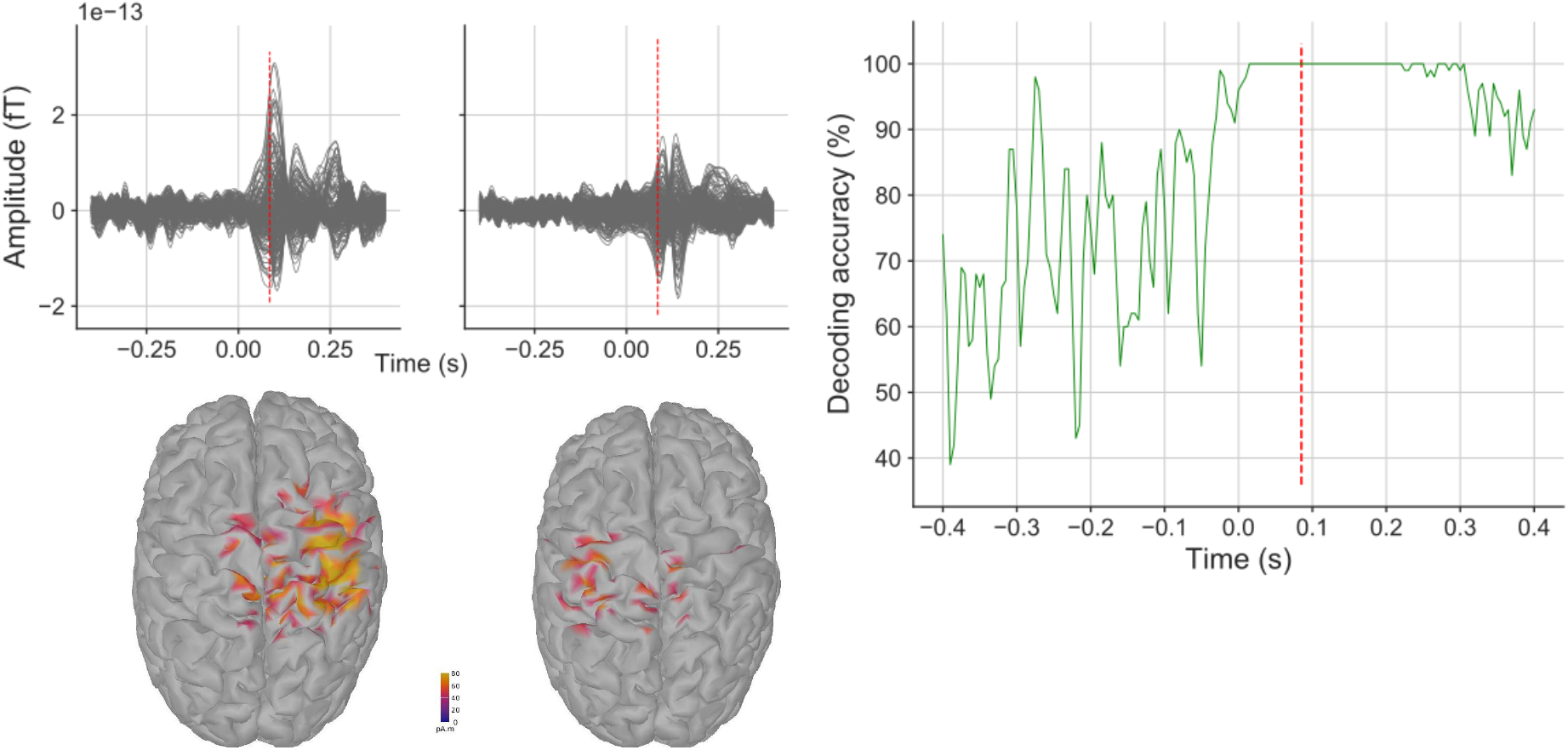
SVM decoding of left versus right button inputs for participant R3154. **The top left two panels** show the averaged 244 channel MEG time series from −400ms to 400ms for left and right translation inputs respectively. **The bottom left two panels** show corresponding heat maps of cortical source estimates in picoAmpere at 0.085 seconds after the input was registered by Meta-T. **The right most panel** shows the averaged percentage accuracy of the classifier at each millisecond of the MEG time series. The red vertical dotted line on each line plot shows the time point corresponding to the presented cortical activity.

### 3.2 Hidden Markov Model analysis

#### 3.2.1 State temporal dynamics

We evaluated our three state HMM using a range of metrics describing both the temporal dynamics of the states as well as Meta-T related performance across each state. The central output of the model is the transition matrix, which describes the probability of participants switching between each pair of states from one tetromino drop to the next. Our transition matrix showed that switches between some pairs of states are more probable than others (Figure 6). In particular, the probability of switches from State 1 to State 1 and State 3 to State 3 were high (0.69 and 0.79 respectively) showing that participants have an affinity to remain in these states once they enter them. The probability of switching from State 1 to State 2 was also relatively high, while the switches from State 2 to State 2 or State 3 were relatively low (0.2 and 0.13 respectively), suggesting that State 2 was a transient state that participants switched to mostly from State 1 but seldom remained in.

**Figure 6:**
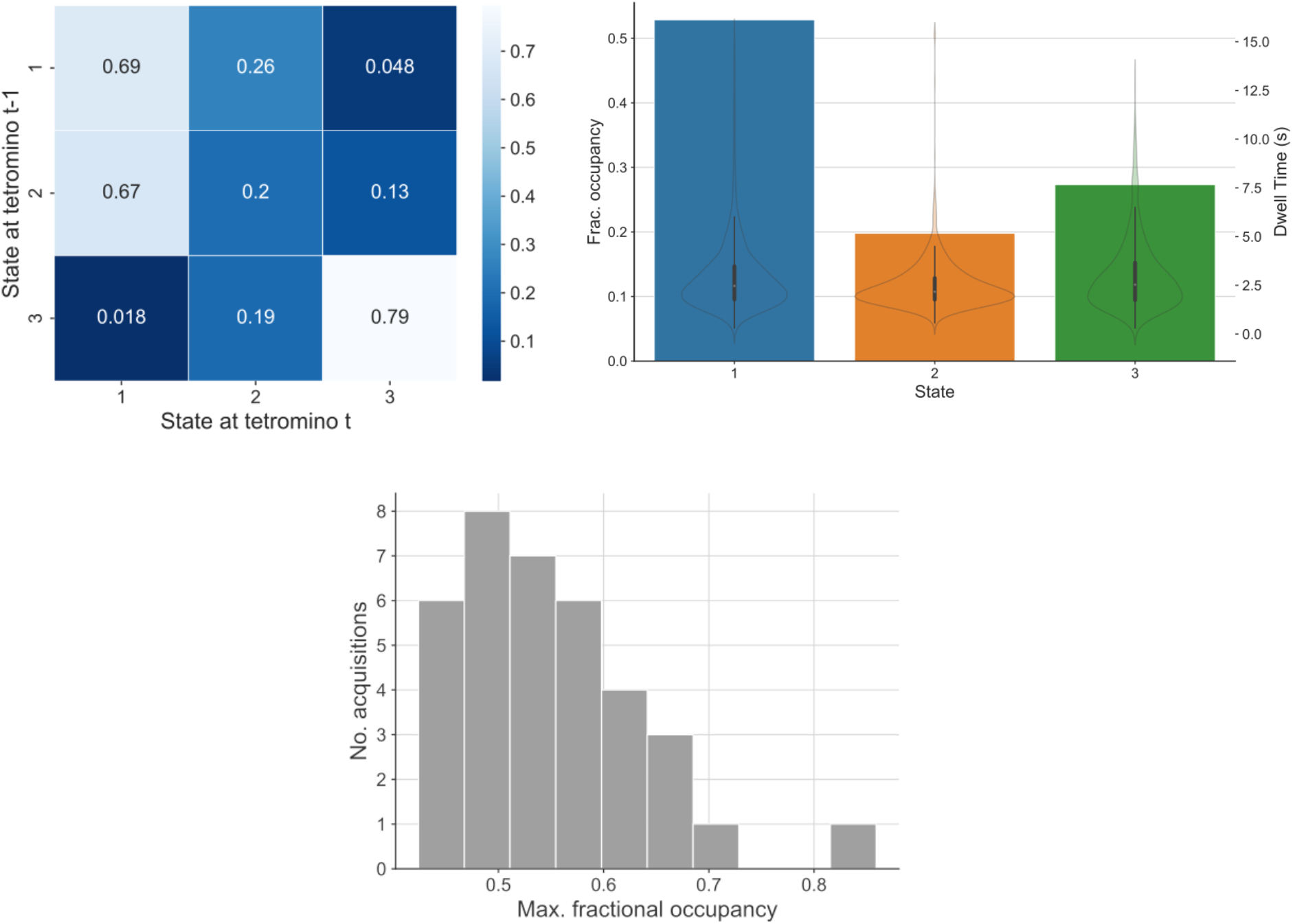
Overview of three state HMM temporal dynamics. **The top left panel** shows the inferred transition matrix of the model, describing the probability of switching between each pair of states. **The top right panel** shows the fractional occupancy for each state, defined as the fraction of total time spent in that state, as well as violin plots depicting distributions of dwell times (the amount of time resided in each state occupancy) across each state. **The bottom panel** shows the distribution of maximum fractional occupancy across acquisitions in the sample. That is, for each data acquisition in the sample, the maximum fractional occupancy represents the fraction of total time spent in the state that the participant occupied for the most amount of time for that acquisition.

To glean further information about the temporal dynamics of our model, we used the Viterbi algorithm to predict the optimal state sequence of our model, and then computed the fractional occupancy of each state, that is, the fraction of total time that is spent by our sample in each state, both in the data set as a whole as well as in each individual game (Figure 6). Previous applications of HMMs to the analysis of human brain dynamics have evaluated HMM validity by examining how state occupancy is distributed across participants. An effective HMM would be expected to output state sequences that show participants occupying multiple states without huge discrepancies in state occupancy (suggestive of single states overwhelming entire participants or recordings). One statistic that reflects this requirement is the maximum fractional occupancy, that is, the fraction of time taken by the state that occupies the most amount of time in a given data acquisition or participant.

To examine this criterion, we visualised our transition matrix together with a bar chart depicting fractional occupancy in each state, as well as a histogram of maximum fractional occupancy across all data acquisitions. In our case, the majority of games had maximum fractional occupancy below 0.6 (mean fractional occupancy was 0.54), demonstrating that our participants’ time was shared across all states in our model. Our plot showed that a little over half of all time (52%) on task was spent in State 1, making this the dominant state throughout task performance. This was followed by State 3, accounting for 28% of state occupancy, and State 2 with 20%.

#### 3.2.2 State performance dynamics

Together these visualisations inform us about how participants transition between and how frequently they occupy states as they play Meta-T, but they do not tell us how behaviour and cognition varies across states. To investigate the dynamics of our performance components across states we started by visualising the time series of observed performance components for individual participants and games in parallel to the time series of posterior probabilities; a secondary output of our model that describes the probability of each of the three states being active given our observations for any given participant and game (Figure 7). By plotting these two time series in parallel, it becomes possible to visually relate patterns of performance to particular states in any given segment of our data. For instance, looking at a game from participant R3154, it is apparent that when well preparation is high, pile disarray is reduced. This pattern appears when the participant enters State 2 which, consistent with our interpretation of the transition matrix, appears to be a transient state with relatively short dwell times. The inverse is the case during State 3, which is associated with low well preparation and increases in pile disarray. It is also apparent that the participant’s motor executions are most efficient during State 1, as is evident from dips in action inefficiency following transitions to this state.

**Figure 7:**
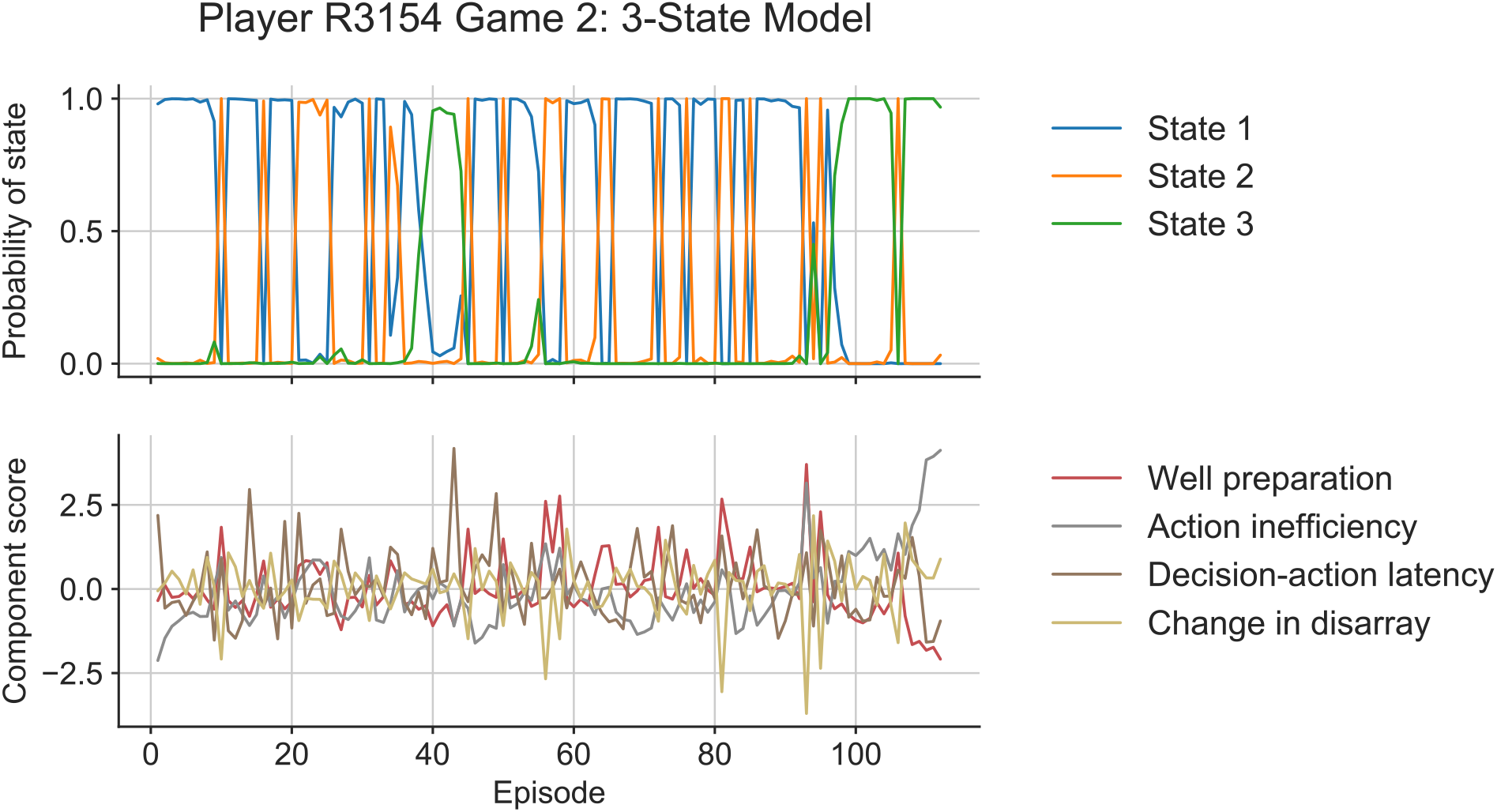
Time series of posterior probabilities and observed performance in an example game. **The top panel** shows the time series of state posterior probabilities, describing the inferred probability of each state being active given the data, across all tetromino drops. **The bottom panel** shows the time series of all four performance components, displayed as z-scores, across all tetromino drops.

#### 3.2.3 State 1

Interpreting performance dynamics across states by visualising individual matches is helpful but not entirely straightforward. To assist in the interpretation of state-performance dynamics across the entire sample, we referred to the mean of each component across each state, which is the primary Gaussian emission returned by our model (see Table X for an overview of these values). We also visualised the distributions of our component scores across each of our states (Figure 8) to an overview of how behaviour varies on average across the three states.

**Figure 8:**
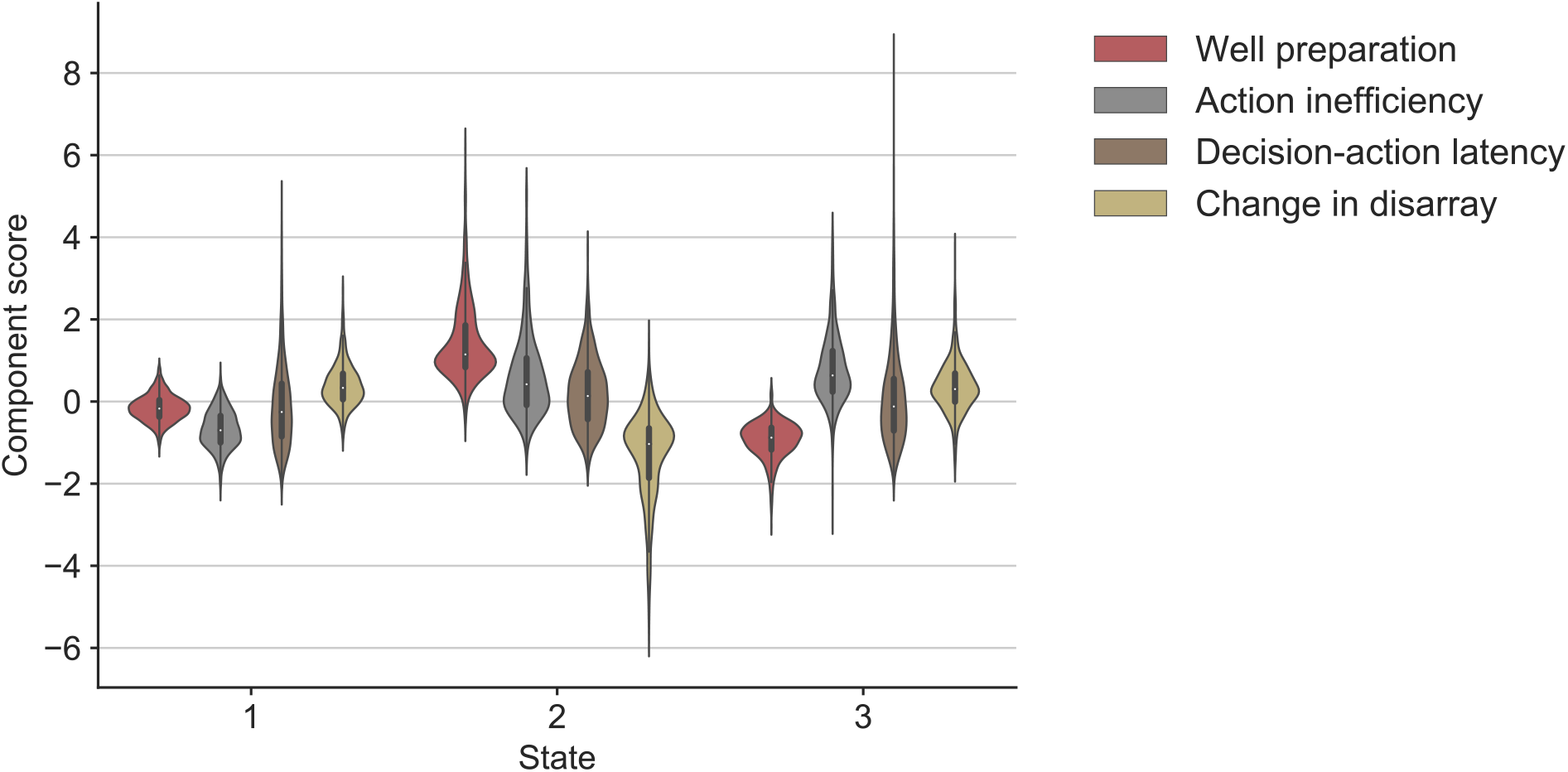
Violin plot of performance component distributions across states. Each “violin” depicts the distribution of the corresponding performance component across all tetromino drops within occurrences of a given state. The black bar in the center of the violin is a box plot, with the center showing the median observed value in the distribution. The coloured portion of the violin shows the kernel density estimate of the distribution. The tails of the violin extending beyond the inner box plot show the range of extreme outliers.

On average, State 1 is characterised by relatively quick decision-making and efficient motor execution, as well as slightly under-average well preparation and slight increases to pile disarray at each tetromino drop. In line with State 1 being the most occupied state across the data set, we view State 1 as the default “engaged” state, corresponding to usual, attentive Tetris gameplay.

**Table 1:**
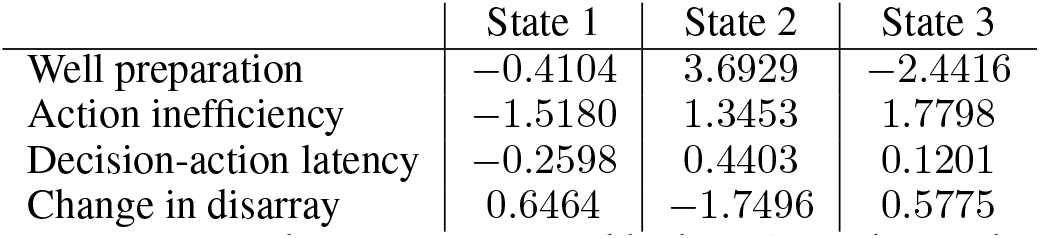
Mean component scores across each state as computed by hmmlearn during the Markov modelling process. Each value describes the mean of the distribution of component scores estimated for the corresponding state.

|  | State 1 | State 2 | State 3 |
| --- | --- | --- | --- |
| Well preparation | -0.4104 | 3.6929 | -2.4416 |
| Action inefficiency | -1.5180 | 1.3453 | 1.7798 |
| Decision-action latency | -0.2598 | 0.4403 | 0.1201 |
| Change in disarray | 0.6464 | -1.7496 | 0.5775 |

#### 3.2.4 State 2

On the other hand, State 2 is characterised by high well preparation, reductions to pile disarray, and high motor inefficiency and decision-action latency. Additionally, given that dwell times in State 2 appear to be short, we interpreted State 2 as a transient “opportunity” state, during which the participant is prepared to either score significant points through line clears, or fumble and compromise the established pile structure. We pursued this idea by calculating the percentage of tetromino drops in State 2 that resulted in at least one line clear. This number was 97%, confirming our initial intuition. The remaining 3% of State 2 drops that did not result in a line clear were distributed across 11 players in the sample, indicating that this state does not exclusively capture cleared lines, but rather pile structure conducive to line clears that most players in the sample occasionally failed to take advantage of.

#### 3.2.5 State 3

Finally, and in contrast to State 2, State 3 was characterised by the lowest well preparation, increases to pile disarray, as well as relatively high motor inefficiency and high but extremely variable decision-action latency. We interpreted State 3 by considering these trends in tandem with aforementioned temporal dynamics. That is, instances of State 3 showed higher dwell times than State 2, and transitions to State 3 were over twice as likely from State 2 than from State 1. Taken together, we interpreted State 3 as a “hyper-engaged” state characterised by poor motor execution and planning, during which participants attempt to resolve difficult pile structures that likely arise from sudden and significant changes to structure that may occur in State 2.

#### 3.2.6 Endogenous rhythms across states

We investigated whether our states were neurally distinct by comparing the averaged amplitude of activity within frequency bands and regions of interest between our states. Specifically, after computing the fourier transform of bins across each state, we aggregated neural activity within states for each participant by computing the RMS of frequency bands corresponding to endogenous rhythms of interest. Principal among these rhythms was the occipital alpha rhythm, which has previously been linked to attention. We also compared mu band activity in the primary motor cortex across states. Interestingly, V1 RMS alpha and M1 RMS mu were different between the left and right hemispheres (see Figure 10 for distributions of the respective observations). For this reason, we conducted separated statistical analyses for each hemisphere.

We first conducted a one-way repeated measures ANOVA to test for within-participants differences in V1 RMS alpha between states. These tests were significant for both left V1 [F(2, 24) = 3.6317, *p* = 0.0419] and right V1 [F(2, 24) = 4.2665, *p* = 0.0260]. Both tests yielded small effect sizes (*η*^2^ = 0.0024 and *η*^2^ = 0.0046 respectively). We also conducted one-way repeated measures ANOVAs to test for within-participants differences in M1 RMS mu between states. Differences in neither left M1 [F(2, 24) = 0.7357, *p* = 0.4896] nor right M1 [F(2, 24) = 0.8488, *p* = 0.4404] were statistically significant.

Post-hoc differences for within-participants differences in occipital alpha across states showed significant differences in alpha activity in the left primary visual cortex between states 1 and 3 (*p* = 0.0374, Cohen’s *d* = −0.1036), as well as significant differences in the right primary visual cortex between states 1 and 2 (*p* = 0.0194, Cohen’s *d* = 0.0835) and states 2 and 3 (*p* = 0.0436, Cohen’s *d* = −0.1585). These results suggest that, in addition to our states displaying distinct patterns of alpha activity, participants manifest the highest levels of alpha in State 3, followed by State 1.

## 4 Discussion

Drawing on recent advances in behavioural neuroscience, we used an HMM to identify hidden states in multivariate psychomotor data obtained from an ecologically valid task, showing that humans shift between latent states during psychomotor performance that differ in behavioural and neural characteristics. Our task was a laboratory version of Tetris that logs granular performance metrics through time, and was performed in an MEG scanner. We identified three states with unique temporal and behavioural dynamics: 1) a default “engaged” state in which participants spent the most amount of time, characterised by averagely-difficult structure of the Tetris pile, quick decision-making and efficient motor execution, 2) an “opportunity” state with the lowest occupancy and shortest dwell times, characterised by points scoring and high well preparation, but poor motor execution and decision-making speed, and 3) a “hyper-engaged” state with the second highest fractional occupancy, where participants contended with difficult pile structures, often with poor motor execution and decision-making speed. In addition to highlighting differences in performance, we showed that states differ in their neural signatures by comparing the amplitudes of endogenous rhythms between states. Comparisons of neural activity between our three states revealed statistically significant differences in amplitudes of occipital alpha-band activity, a signal associated with attentional state, indicating that differences in cognition acros states may relate to attention. Taken together, our findings show that humans switch between behaviourally and neurally distinct states as they engage in complex psychomotor performance. We show that the dynamics of these state transitions can be captured using synchronised behavioural and neural measurements, and subsequently modelled using unsupervised learning techniques to describe the relationship between latent states and performance.

Previous latent state models of behaviour have concentrated on animal behaviour in relatively well-studied task environments, such as courtship behaviours in fruit flies [17], visual detection in mice [16, 26, 18], or swim bouts in larval zebrafish [62]. These paradigms lend themselves well to models of latent states as the resultant observations are intuitively discretisable. Additionally, many of these studies are high in ecological validity, modelling behaviours that would be natural to observe in an animal’s usual behavioural repertoire. In comparison, the application of sequence classification techniques to identify latent states in humans has predominantly involved artificial tasks (e.g., motion coherence task [18]) or resting-state FMRI [27, 23, 29]. In the present work, we used a laboratory adaptation of a highly popular and commercially successful video game, paralleling growing interest in the use of naturalistic stimuli within the domain of cognitive neuroscience [63, 64]. Specifically, participants played a laboratory adaptation of Tetris [50, 19] that collects numerous cognitive-behavioural variables relating to game state, motor execution, and motor planning. Our analysis included a feature engineering component whereby behavioural measurements were decomposed into four performance components based on data obtained by an independent laboratory using the same task [53]. Thus, using a tried and tested version of a video game explicitly tailored for laboratory research, we add to a growing body of literature that uses video games for research in cognitive neuroscience [7, 65, 66, 67].

Many studies of cognition that use video games, in particular commercial video games, analyse univariate measures of performance such as end of match summary metrics (e.g., win/loss, points scored), or time-bound measures of performance (e.g., points scored per minute). We show here that the analysis of multivariate behavioural time series can generate inferences and research questions that may be difficult to access with summary metrics alone. Relatedly, and partly as a consequence of this limitation, studies of video games that involve repeated measures often aggregate data within and across sessions of engagement. Previous work has advised against this on theoretical [11, 12, 13] as well as empirical grounds, demonstrating how certain insights into individual differences [21] or skill acquisition [68, 69] can only be achieved after disaggregating data and considering behaviour in a more detailed fashion. Although this study involved detailed analysis of behaviour through time, we are guilty of the sin of aggregation as we too considered our sample as a single homogenous group, despite variation in players’ average scores indicating a heterogeneity in skill level.

One related implication for our analysis is that phases of gameplay that are more demanding for less skilled players may place lower demands on best players in the sample. Having concatenated all observations to produce our input time series for model fitting, our model would not have accounted for the potential effects of variation in skill. This is an important consideration, given previous evidence highlighting that variables discriminating between less versus more skilled players are not the same across skill brackets [70]. In parallel research involving animal subjects, this issue is either resolved through extensive training, or it is completely bypassed by observing naturally ingrained behaviours. Presently, we made efforts to recruit participants who reported familiarity with Tetris, but we were unable to control for how proficient they were. Additionally, we realised during data collection that many participants were familiar with modern versions of Tetris with nuanced differences that confounded their initial experiences for the game. For instance, our configuration of Meta-T emulates Classic Tetris and therefore lacks visual guidelines indicating each tetromino’s destination, and prohibits rotating tetrominoes at the very edge of the well; both of these are mechanics that some of our more experienced participants reported relying on in their usual recreational gameplay. We acknowledge that these issues are likely to have introduced noise to our model.

Compared to previous latent state models of low-level psychophysical phenomena, we opted for a complex behavioural environment that is high in ecological validity. In doing so, we show that hidden Markov modelling can be used to identify state shifts in tasks that approach real world behaviour. However, we acknowledge our position in the trade-off between simple behavioural data suitable for predictive modelling versus rich behavioural data that makes prediction much more difficult. Given the nature of our input data (i.e., our time series of performance components), our model infers parameters that describe the temporal dynamics of our states, and generates emission probabilities describing the probabilities of observations given the state time series. In the case of our Gaussian HMM, the emission probability parameters of each state were the mean and standard deviation parameters describing the Gaussian probability density function of each performance component in the respective state. Consequently, this can lead to expectations about how participants may perform across states based on how the distributions of performance components shift across states. However, this does not permit the more fine-grained predictions that other studies have done, such as the GLM-HMM approach in [17].

The validity of our model is supported somewhat by the correspondence between the behavioural characterisations of our states and the underlying neural signatures of each state. That is, in a model that failed to distinguish between cognitively meaningful states, we would expect to observe no differences in neural signatures associated with cognition. Instead, comparisons of neural activity across our inferred states revealed statistically significant differences in occipital alpha, a signal that has been previously linked to attention. In particular, post-hoc tests revealed elevated occipital alpha in State 3 as compared to State 1, and higher occipital alpha in State 1 as compared to State 2. Given the existing association between occipital alpha and attention, this result is consistent with our analysis of performance, which was suggestive of increased attentional demand in State 3 due to the presence of difficult pile structures and large variance in participants’ decision-making latency. However, although it is likely that our model is detecting shifts in attention, it is difficult to infer precisely what aspects of the task are being attended to across different states, as we did not manipulate attention as previous experiments investigating performance and attention have done [71, 72].

It is also possible that attention shifts continuously, and not discretely as assumed by our model. Ashwood and colleagues ([18]) found superior model fit in their discrete model as compared to a model with continuous latent states [26], albeit in the context of a different task. Additionally, these authors found that a two-state discrete model fit human data from a motion coherence task better than a three-state model. We are open to the possibility that models with different assumptions may describe performance in the present context better than our three-state guassian HMM, for instance a two-state, engaged versus disengaged model. However, this is a question for future work.

### 4.1 Limitations

A limitation of our study is that, in contrast to previous latent state models of behaviour, we did not have adequate time to train our participants on the task. Ours was a complex psychomotor task requiring both rapid perceptual decision-making and skilled motor inputs. Although we recruited participants who all indicated ample prior experience with Tetris, as mentioned before, we are nonetheless conscious of large variation in participant skill, as well as noise arising from unfamiliarity with our specific configuration of Meta-T, the controller, and the scanner environment in which the task was performed. In addition, we note the absence of a “ground truth” model with which to validate our model. Instead, we compared the log-likelihood of our model to a randomised chance model, which indicated superior fit of the true model. However, we acknowledge as a limitation that due to the nature of our input data and the type of HMM that we used, the predictive capacity of our model is restricted.

### 4.2 Conclusion

Studies that take repeated measurements of performance in a task, such as episodic performance in a video game, often average data within and across sessions. Recent work has demonstrated that, in the context of repeated trials within sessions of behaviour, animals and humans shift between discrete behavioural strategies during performance. Using simultaneous behavioural and neural recordings of participants playing a laboratory version of Tetris, we extend previous work by demonstrating that individuals switch between latent states during performance in an ecologically valid task. Individuals in our sample shifted between three states each with unique performance characteristics during gameplay. Further, MEG analysis revealed differences in occipital alpha across states, suggesting that differences across states may be related to attention. Our results show that analysing sessions of data by averaging summary statistics alone may mask a wealth of information describing the dynamics of performance and cognition. We demonstrate how these dynamics can be uncovered unsupervised learning techniques and granular, multivariate data.

## A Appendix

**Figure 9:**
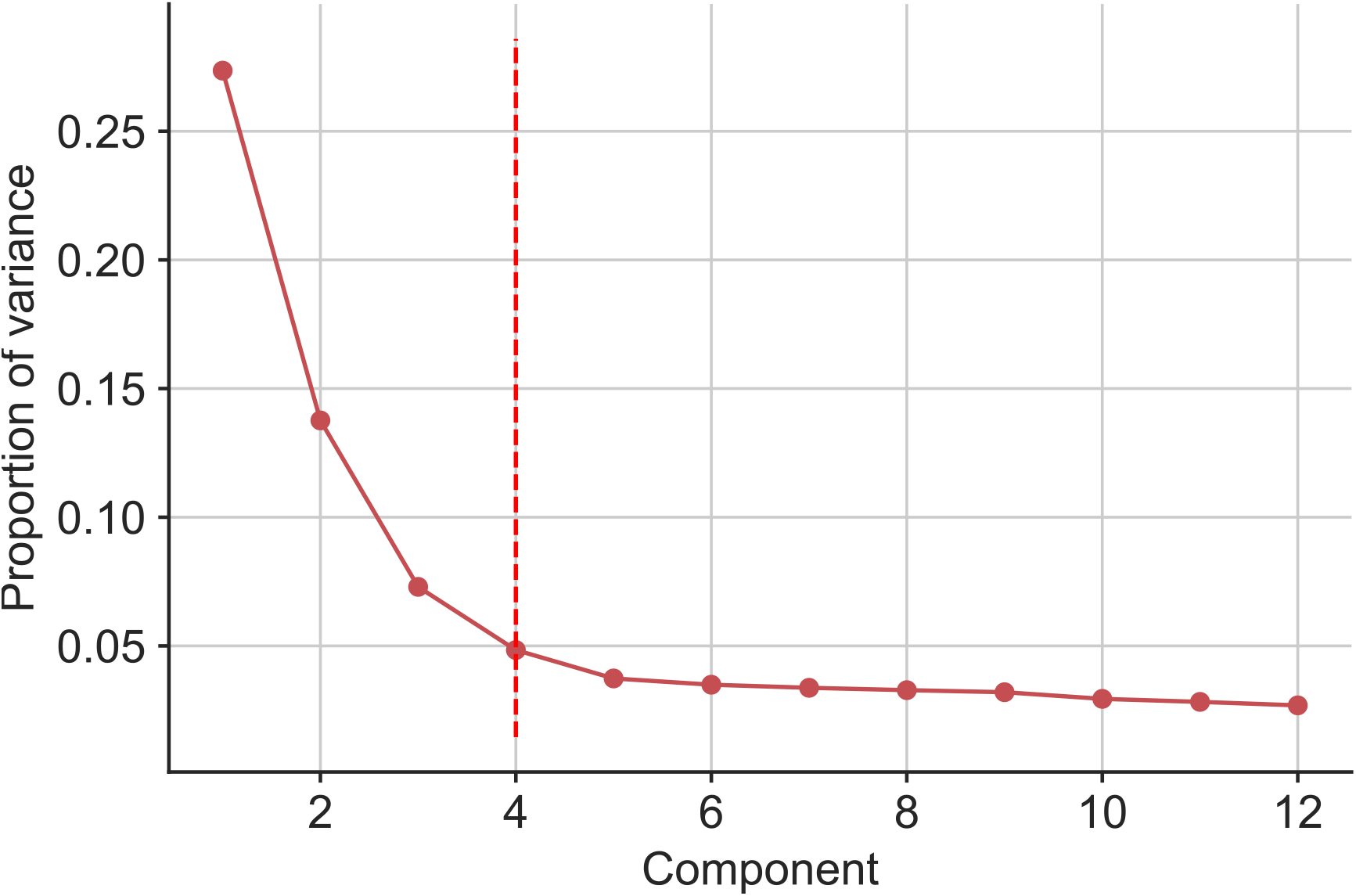
Scree plot of PCA. Points in the line show the variance explained by each successive principal component, where components are ordered from most to least variance explained. The vertical dashed line indicates the point of maximum curvature in the line, corresponding to our choice of number of components to be generated for our analysis.

**Table 2:** Table of PCA loadings. Column headers display the number of each component ordered by descending amount of variance explained. Cells show the correlations past 0.20 between variables (labelled in the row headers) and the respective components.

|  | 1 | 2 | 3 | 4 |
| --- | --- | --- | --- | --- |
| mean_ht | 0.2960 |  |  |  |
| row_trans | 0.2890 |  |  |  |
| weighted_cells | 0.2890 |  |  |  |
| max_ht | 0.2870 |  |  |  |
| landing_height | 0.2850 |  |  |  |
| pit_rows | 0.2790 |  |  |  |
| pit_depth | 0.2730 |  |  |  |
| col_trans | 0.2710 |  |  |  |
| pits | 0.2710 |  |  |  |
| lumped_pits | 0.2670 |  |  |  |
| min_ht | 0.2530 | -0.2080 |  |  |
| wells |  | 0.3810 |  |  |
| deep_wells |  | 0.3800 |  |  |
| max_well |  | 0.3720 |  |  |
| cuml_wells |  | 0.3690 |  |  |
| jaggedness |  | 0.3220 |  |  |
| max_diffs |  | 0.2770 |  |  |
| cd_1 |  | 0.2110 |  |  |
| cd_9 |  | -0.2090 |  |  |
| min_rots_diff |  |  | 0.4440 |  |
| rots |  |  | 0.4420 | -0.2230 |
| min_trans_diff |  |  | 0.3980 |  |
| drop_lat |  |  | 0.3730 |  |
| trans |  |  | 0.3670 | -0.2460 |
| prop_u_drops |  |  | -0.2590 |  |
| avg_lat |  |  |  | 0.5070 |
| initial_lat |  |  |  | 0.4560 |
| d_max_ht |  |  |  | 0.3550 |
| matches |  |  |  | -0.3690 |
| cd_2 |  |  |  |  |
| cd_3 |  |  |  |  |
| cd_4 |  |  |  |  |
| cd_5 |  |  |  |  |
| cd_6 |  |  |  |  |
| cd_7 |  |  |  |  |
| cd_8 |  |  |  |  |
| pattern_div |  |  |  |  |
| resp_lat |  |  |  |  |

**Figure 10:**
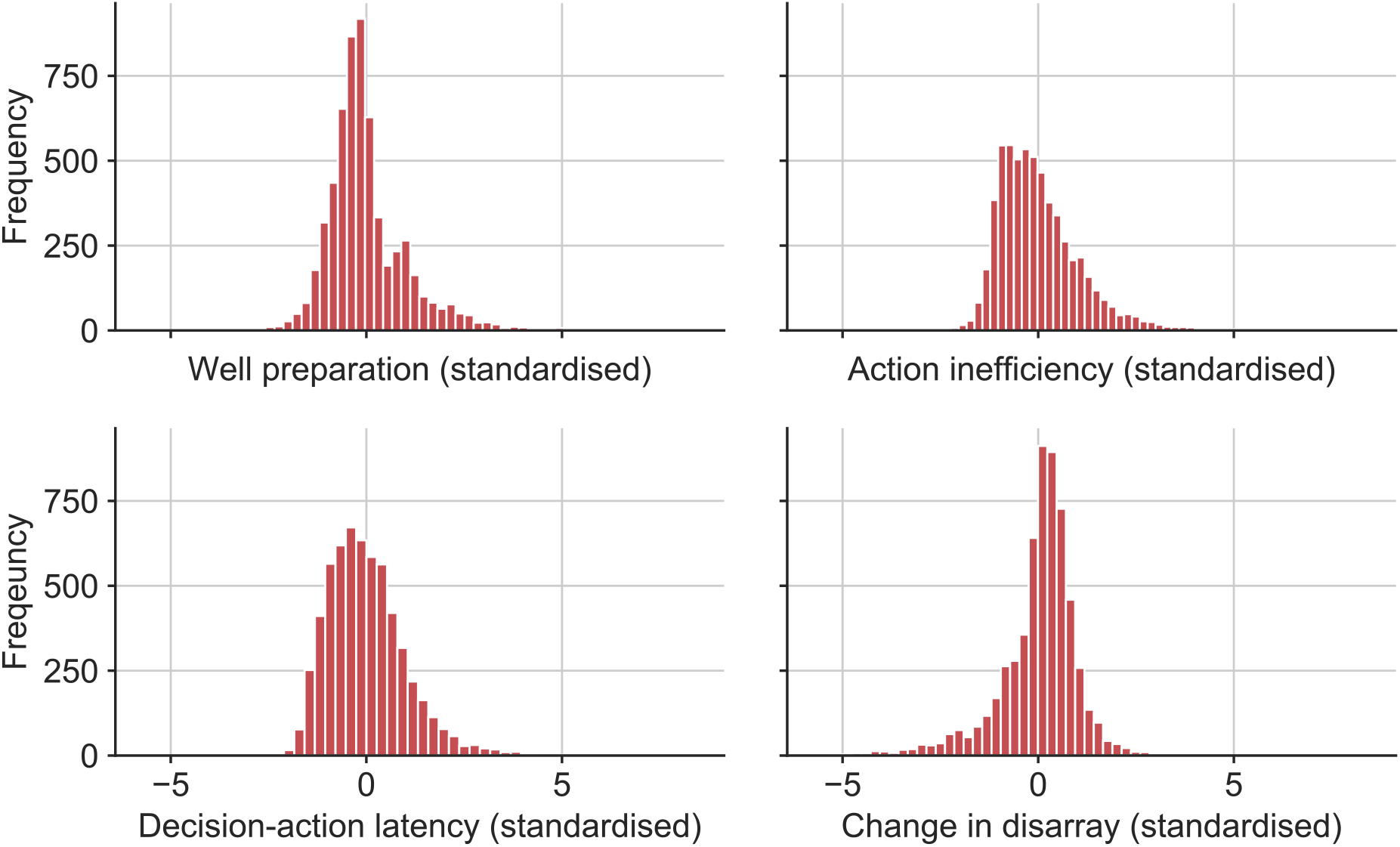
Distributions of component score at each tetromino drop for each component. From top left going clockwise to bottom left, the plots show histograms of z-scored well preparation, action inefficiency, decision-action latency, and change in disarray respectively

